# Isolating mitotic and meiotic germ cells from male mice by developmental synchronization, staging, and sorting

**DOI:** 10.1101/342832

**Authors:** Katherine A. Romer, Dirk G. de Rooij, David C. Page

**Author notes:** Corresponding author at: Whitehead Institute, 455 Main Street, Cambridge, MA 02142, USA. *Email address:* (D. Page).

## Abstract

Isolating discrete populations of germ cells from the mouse testis is challenging, because the adult testis contains germ cells at every step of spermatogenesis, in addition to somatic cells. We present a novel method for isolating precise, high-purity populations of male germ cells. We first synchronize germ cell development in vivo by manipulating retinoic acid metabolism, and perform histological staging to verify synchronization. We use fluorescence-activated cell sorting to separate the synchronized differentiating germ cells from contaminating somatic and germline stem cells. We achieve ∼90% purity at each step of development from the germline stem cell pool through late meiotic prophase. Utilizing this “3S” method (synchronize, stage, and sort), we can separate germ cell types that were previously challenging or impossible to distinguish, with sufficient yield for epigenetic and biochemical studies. The 3S method should enable detailed characterization of molecular changes that occur during the mitotic and meiotic phases of spermatogenesis.

## INTRODUCTION

Surprisingly little is known about the molecular and biological changes that occur during mammalian spermatogenesis—the transformation of stem cells into highly specialized spermatozoa. During this process, germ cells proliferate mitotically and lose pluripotency, and then divide meiotically to produce haploid gametes, which undergo further differentiation to become mature spermatozoa. Understanding the precise molecular and biochemical changes underlying spermatogenesis would require isolating each step along this developmental progression – an extraordinarily difficult effort because the adult testis contains both somatic cells and millions of asynchronously developing germ cells. We present here a novel method, “3S” (synchronize, stage, and sort), which isolates precise populations of mitotic and meiotic germ cells for study, circumventing the extreme cellular complexity of the testis. 3S combines two approaches to germ cell isolation—in vivo simplification of the cellular complement of the testis, followed by ex vivo cell sorting.

Previous attempts to isolate specific germ cell populations have largely relied on cell sorting, using the unperturbed wild-type testis as starting material. Unfortunately, specific molecular markers of most steps of germ cell development have not yet been identified, with the notable exception of the germline stem cell pool (Oatley and Brinster, 2008; Shinohara et al., 2000). Thus, investigators have relied on more general cell properties to separate different germ cell populations. Sedimentation-based approaches, including unit gravity sedimentation and centrifugal elutriation, separate germ cells based on size and density (Bellvé, 1993; Grabske et al., 1975). DNA staining/FACS (fluorescence-activated cell sorting) uses a dye such as Hoechst 33342 to separate germ cells based on DNA staining and light scattering properties (Bastos et al., 2005; Gaysinskaya et al., 2014). Both of these approaches require substantial technical expertise, and both are powerful when used correctly: they can, with high accuracy, separate mitotic, meiotic, and post-meiotic germ cells from one another and from the somatic cells of the testis. However, both approaches have limited resolution: they frequently combine multiple closely related steps of germ cell development into a single sorted population. Resolution is especially limited in the mitotic and early meiotic phases of spermatogenesis, because these cell populations make up a relatively small fraction of the adult testis (Gaysinskaya et al., 2014).

To gain a detailed understanding of the molecular events of spermatogenesis, we need to be able to accurately separate closely related, low-abundance germ cell subpopulations. For example, the mitotic phase encompasses seven histologically distinct steps of germ cell development (A_undiff_/A_1_/A_2_/A_3_/A_4_/Intermediate/B spermatogonia, Fig. 1A), which DNA staining/FACS cannot separate, and which unit gravity sedimentation can separate into only two populations. During this mitotic phase, germ cells undergo important functional changes, losing expression of pluripotency markers and acquiring competence to enter meiosis (Buaas et al., 2004; Endo et al., 2015; Shinohara et al., 2000). However, the gene expression changes that underlie these functional changes are still poorly understood (de Rooij and Griswold, 2012; Zhang et al., 2014). Similarly, the first two steps of meiotic prophase, leptotene and zygotene, are histologically distinct and encompass distinct chromosomal events, with programmed double-strand breaks occurring during leptotene, and synapsis of homologous chromosomes beginning at zygotene (Fig. 1A). These populations cannot be separated with sedimentation-based approaches, and are difficult to separate with DNA staining/FACS approaches (Bellvé, 1993; Gaysinskaya et al., 2014). If leptotene and zygotene germ cells could be effectively separated, gene expression differences and chromatin state could be systematically characterized (da Cruz et al., 2016; Gaysinskaya et al., 2014).

**Fig. 1.**
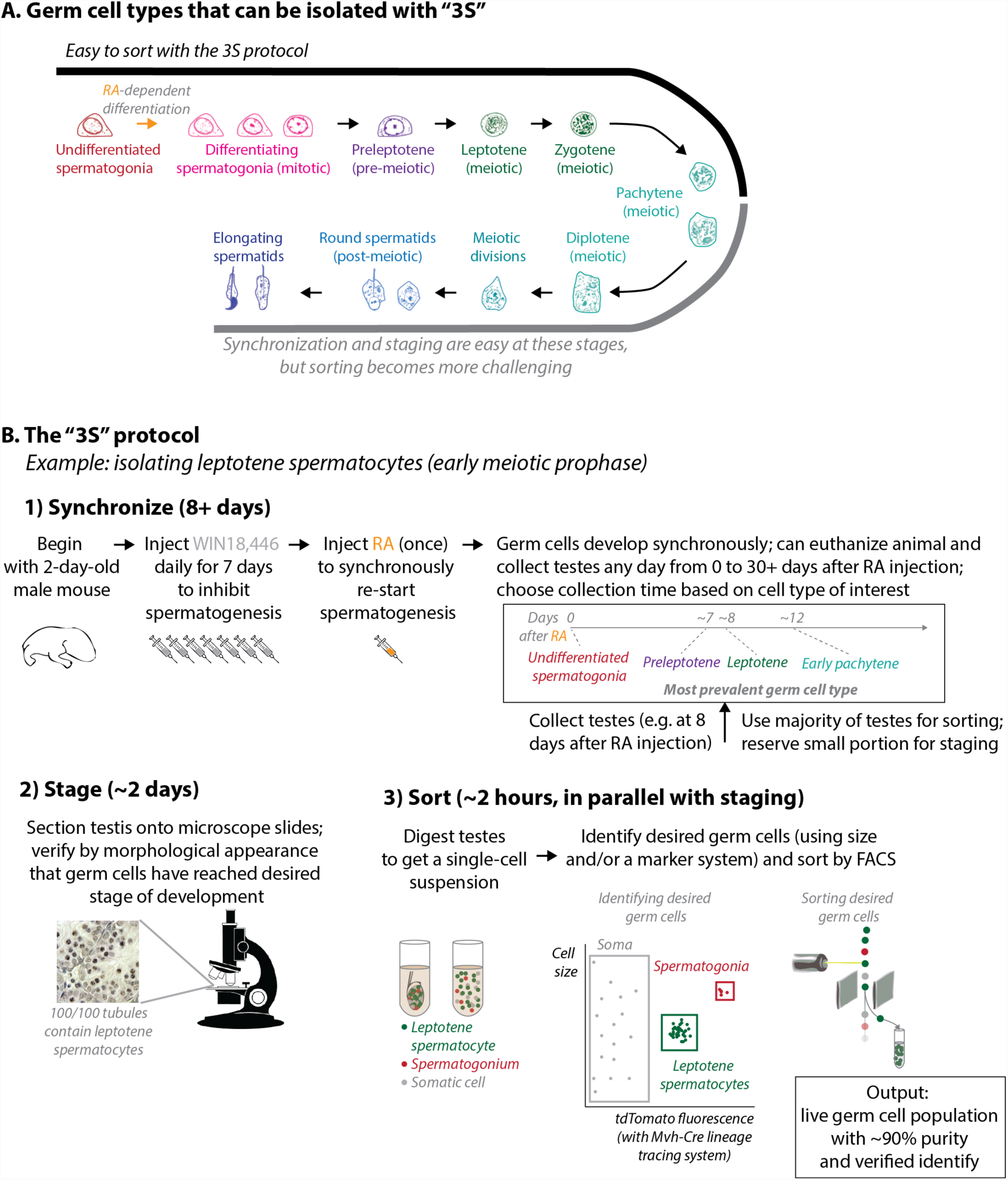
Overview of “3S” (synchronize, sort, and stage) (A) Schematic of germ cell development in the testis of an unperturbed male mouse, showing germ cell populations that can be isolated with “3S.” In the neonatal mouse, the only germ cells present are *undifferentiated spermatogonia*: these encompass the stem cells, which persist throughout the animal’s lifetime. In a retinoic acid (RA)-dependent fashion, cohorts of these undifferentiated spermatogonia then differentiate to become mitotic *differentiating spermatogonia* (A_1_, A_2_, A_3_, A_4_, Intermediate, and B). Germ cells next initiate meiosis and undergo pre-meiotic DNA replication as *preleptotene spermatocytes*, and then go through meiotic prophase (*leptotene, zygotene, pachytene* and *diplotene spermatocytes*) and two reductive divisions. Finally, germ cells go through post-meiotic development (*round* and *elongating spermatids*). Populations from undifferentiated spermatogonia through mid-pachytene (black bar) can be easily isolated with 3S using the Mvh-Cre/tdTomato germ cell lineage tracing system (Fig. 6). Later steps of germ cell development (gray bar) are more difficult to isolate by sorting, due both to differentiation of subsequent cohorts of undifferentiated spermatogonia and to diminished fluorescence from the Mvh-Cre/tdTomato lineage tracing system. (B) The “3S” protocol. B-1: Description of the synchronization procedure. WIN18,446 (gray) inhibits RA synthesis. The provided timeline for synchronous development after RA injection is for a typical C57BL/6 mouse. B-2: Description of the staging procedure. A small portion of the synchronized testis is embedded in paraffin, sectioned, and examined with a light microscope, to confirm that germ cells have reached the desired step of development. Staging verifies the identity of the sorted germ cells. B-3: Description of the sorting procedure (performed in parallel with staging). Schematic shows digestion of the testis into a single-cell suspension, followed by FACS (fluorescence-activated cell sorting). Schematic for FACS shows identification of three cell populations (leptotene spermatocytes, spermatogonia, and somatic cells) by a combination of cell size, as approximated by light scatter, and tdTomato fluorescence intensity. Based on this identification, leptotene spermatocytes are selectively sorted. For a description of the Mvh-Cre/tdTomato germ cell lineage tracing system, see Fig. 6.

When separating germ cell subpopulations, a common approach to avoiding the complexity of the adult testis is to use the juvenile testis, which displays more circumscribed cellular heterogeneity. Since spermatogenesis begins after birth, one can isolate testes that are enriched for mitotic or meiotic germ cells by timed collection (Bellvé, 1993; Margolin et al., 2014). However, asynchrony is present from the very beginning of spermatogenesis in neonatal males, so sorting precise germ cell subpopulations from the juvenile testis is still challenging (Nebel et al., 1961; Snyder et al., 2010). A more promising approach is to induce synchronization of germ cell development by manipulating retinoic acid (RA) levels in vivo. In the absence of RA, germ cells arrest as “undifferentiated spermatogonia,” a population which encompasses the germline stem cells. A single pulse of RA can then induce spermatogonial differentiation throughout the entire testis. Spermatogenesis proceeds synchronously for months thereafter.

Synchronization was first achieved by withholding and then restoring vitamin A, the metabolic precursor of RA, in the diets of adult mice (Morales and Griswold, 1987; van Pelt and de Rooij, 1990). However, synchronization by vitamin A deprivation is not practical to apply on a large scale: it is time-consuming, requiring approximately five months of dietary manipulation and more than two months of daily animal weighing, and it is quite detrimental to animal health. More recently, Hogarth et al. (2013) developed a practical synchronization method using the compound WIN18,446 [also known as N,N′-1,8-Octanediylbis(2,2-dichloroacetamide)], which inhibits RA synthesis. In these studies, Hogarth et al. administered WIN18,446 to juvenile mice for 7 days (d), followed by a single dose of RA; they achieved synchronization with only 8 d of animal manipulation. Following either of these synchronization protocols, the cell complement of the testis is dramatically simplified: for instance, a synchronized testis might contain germ cells in leptotene, but not germ cells in any other phase of meiosis (Fig. 1B).

Our 3S (synchronize, stage, and sort) method combines these two complementary approaches: in vivo simplification of the cellular complement of the testis, and ex vivo cell sorting. First, we have developed a refined WIN18,446/RA synchronization protocol. This protocol yields predictable timing of germ cell development, which allows us to enrich for precise steps of development simply by timed collection of the testis (Fig. 1B-1). Second, we have developed FACS protocols to efficiently sort the synchronized germ cells from contaminating somatic and undifferentiated spermatogonial populations (Fig 1B-3). Histological staging of a reserved portion of the synchronized testis allows us to verify that synchronization was successful and that we have isolated the desired cell population (Fig. 1B-2). With 3S, we have achieved ∼90% purity of germ cell subpopulations from the undifferentiated spermatogonia through late meiotic prophase and beyond.

## RESULTS

We present in turn detailed protocols and typical results for synchronization, staging, and sorting of male germ cells.

### Step 1: Synchronization

As shown in Fig. 1B-1, we first synchronize germ cell development *in vivo*, such that, at the time of collection, the testis is highly enriched for the germ cell population of interest. Synchronization requires a series of eight daily injections: seven injections of WIN18,446 to hold spermatogonia in an undifferentiated state, followed by a single RA injection to synchronously induce spermatogonial differentiation. After these injections, the synchronized cohort of germ cells follows the developmental progression shown in Fig. 1A. They first go through six mitotic divisions, becoming in turn A_1_, A_2_, A_3_, A_4_, Intermediate, and B spermatogonia. After the sixth mitotic division, the germ cells become preleptotene spermatocytes, during which stage they initiate meiosis and undergo pre-meiotic DNA replication. They then enter meiotic prophase, becoming in turn leptotene, zygotene, pachytene, diplotene, and secondary (dividing) spermatocytes, before undergoing post-meiotic differentiation as round and elongating spermatids (Russell et al., 1990). Based on this developmental progression, we collect tissue at the time-point dictated by the cell population of interest (Figs. 2, 3, 4).

**Fig. 2.**
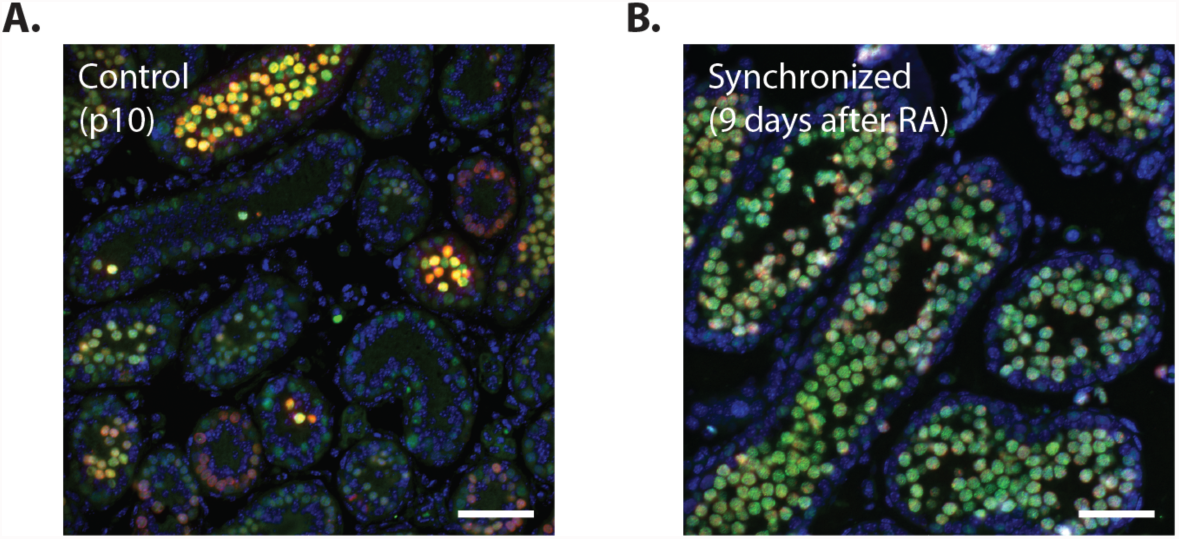
An example of synchronization. Testis sections, stained for SYCP3 (green) and γH2AX (red) and counterstained with DAPI (blue). Cells with high levels of SYCP3 and low/punctate γH2AX are in late leptotene, approaching zygotene. Scale bars, 100 µm. (A) Control, unsynchronized pubertal testis, p10. At this time-point, the unsynchronized testis is relatively enriched for leptotene spermatocytes. Nevertheless, the staining patterns of individual tubules are highly heterogeneous. Most tubules have little or no SYCP3 expression, indicating that they do not contain meiotic spermatocytes. (B) Synchronized testis, 9 d after RA restoration. All tubules are dominated by cells with high SYCP3/low γH2AX, indicative of synchronization at the end of leptotene.

**Fig. 3.**
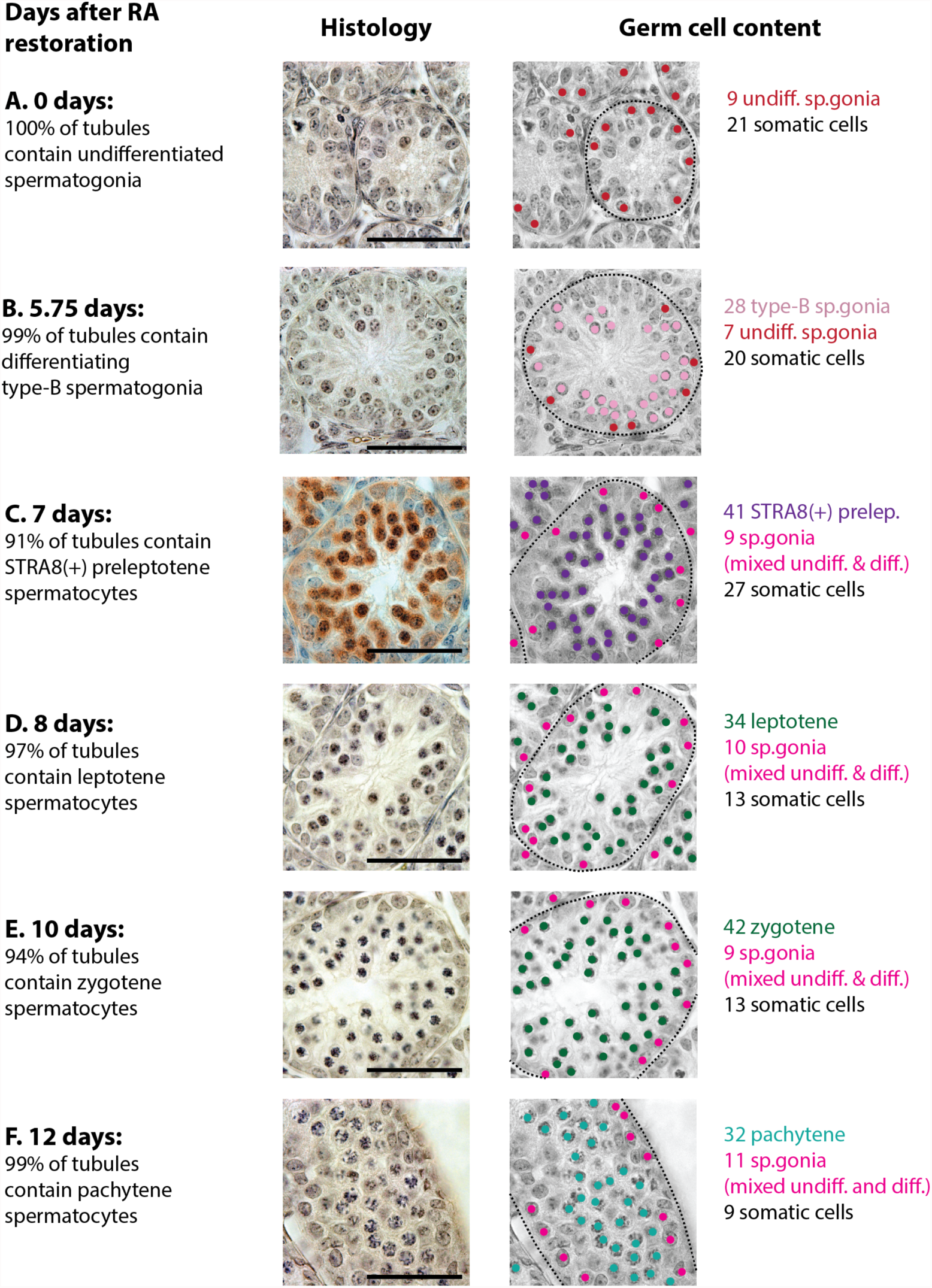
Typical time-course of synchronization. Left column: Days after RA restoration, and most advanced (and most prevalent) steps of germ cell development observed. A minimum of 100 tubules was scored for each sample. Middle column: histology. Section in (C) is stained for STRA8 (brown) and counter-stained with hematoxylin (blue); all other sections are stained with hematoxylin only. Scale bars, 100 µm. Right column: Steps of germ cell development present in each tubule. Grayscale version of image in middle column, with all germ cells in the section marked with colored dots as indicated. All unmarked cells are somatic. For each time-point (A-F), the cells in one example tubule (indicated by dashed black outline) are counted. Undiff.: undifferentiated. Diff.: differentiating. (A) Just before RA restoration. 100% of tubules contain undifferentiated spermatogonia as their only germ cell population. Thus, undifferentiated spermatogonia and somatic cells are the only cell types in the testis; most cells in the testis at this time-point are somatic. (B) 5.75 d after RA restoration. 99% of tubules contain type-B (late differentiating) spermatogonia as their most developmentally advanced (and most prevalent) germ cell population. Tubules also contain somatic cells and a few undifferentiated spermatogonia. These three cell types make up the great bulk of this testis. The 1% of tubules without type-B spermatogonia, which arise due to imperfect synchronization, contribute a few Intermediate spermatogonia. (C, D, E, and F). 7, 8, 10, and 12 d after RA restoration. 91%, 97%, 94%, and 99% of tubules contain, respectively, STRA8-positive preleptotene, leptotene, zygotene, or pachytene spermatocytes as their most developmentally advanced germ cell populations. Tubules also contain somatic cells and a few spermatogonia. The spermatogonia are a mixture of undifferentiated and differentiating spermatogonia, with the undifferentiated spermatogonia encompassing the spermatogonial stem cells, and the differentiating spermatogonia representing a second synchronous generation of germ cells, developing behind the meiotic cohort.

**Fig. 4.**
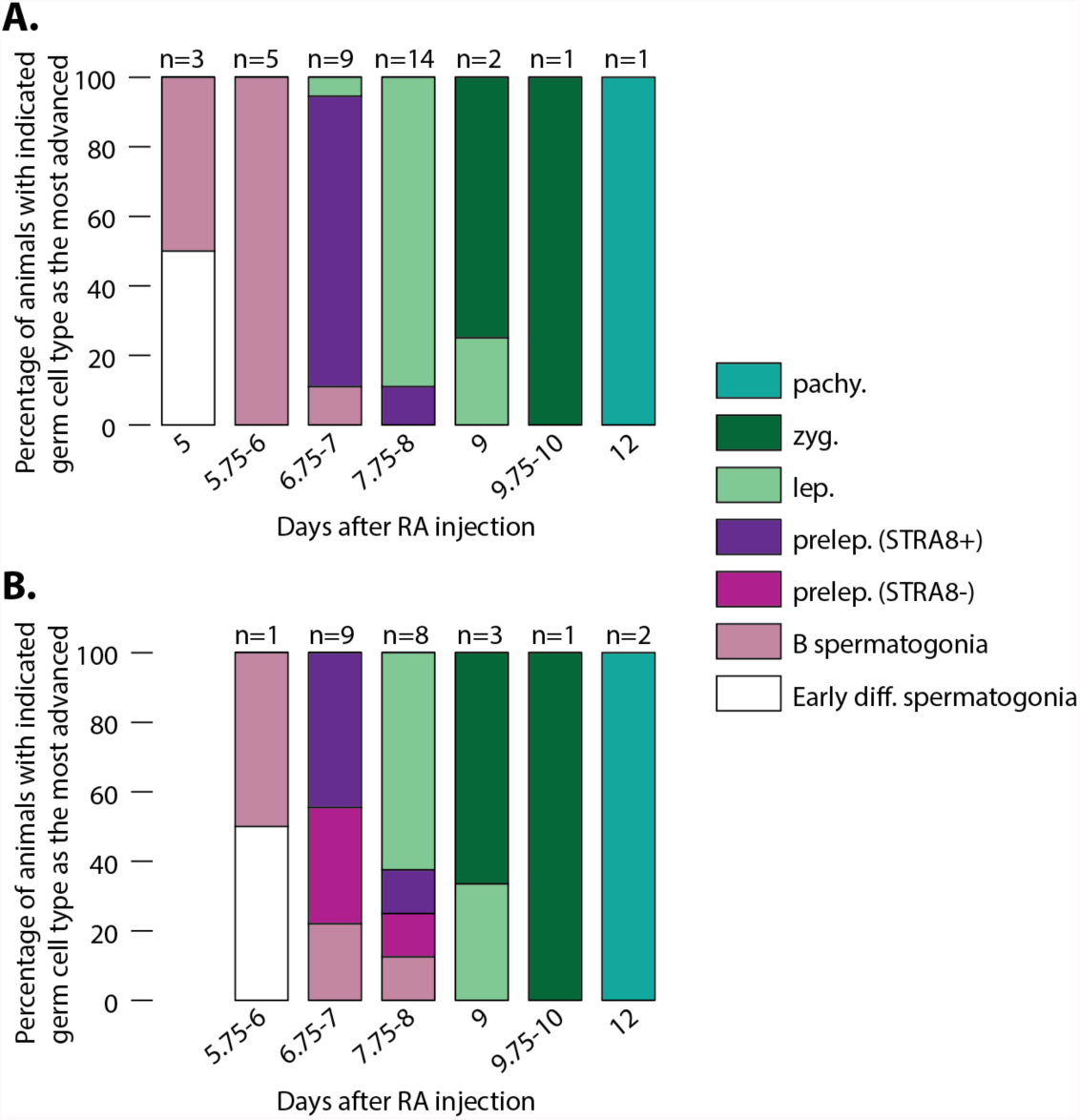
Reproducibility of timing of germ cell development after synchronization. (A and B) Timing of germ cell development in a total of 35 (A) and 24 (B) developmentally synchronized C57BL/6 mice. All of the animals in (A) were of normal weight (**>**5 g at the time of RA injection); all of the animals in (B) were underweight (<5 g at the time of RA injection). For each animal, we staged at least 100 tubules (i.e., determined the most advanced step of germ cell development present in each tubule, as in Fig. 3). If >90% of tubules displayed the same stage, the animal’s count was assigned to that stage. Otherwise, the animal’s count was split equally across all stages that were observed in >10% of tubules. Diff.: differentiating. B: type-B. Prelep., lep., zyg., pachy.: preleptotene, leptotene, zygotene and pachytene spermatocytes. In the normal-weight animals (A), timing of germ cell development after RA restoration is quite consistent, following the schedule presented in Fig. 3. In underweight animals (B), germ cell development is delayed, and its timing is less predictable.

We now describe the protocol for synchronization in detail.

*Materials:*

- Mice, 2 days (d) after birth (p2): Any strain of mice can be used, but the time-course of development after synchronization must be established anew in each strain. To increase the reproducibility of developmental timing, we strongly recommend using genetically uniform mice (inbred or F1 of two inbred strains). We use the C57BL/6 strain of mice. To obtain mice at p2, we either perform timed matings, or check daily for pups. To facilitate the sorting of germ cells from soma, we use a genetically encoded lineage tracing system, which consists of two components. First, our mice are heterozygous for “tdTomato”: a loxP-STOP-loxP-tdTomato construct in the *Rosa26* locus, strain B6.Cg-Gt(ROSA)26Sor^tm14(CAG-tdTomato)Hze^/J (Madisen et al., 2010) from The Jackson Laboratory. Second, our mice are heterozygous for “Mvh-Cre”: a Cre-mOrange fusion protein, driven by the endogenous *Mvh* (*Mouse vasa homolog*, a.k.a. *Ddx4*) promoter (Hu et al., 2013). (We note that the mOrange fluorescence is too weak to use for sorting.) These together form the “Mvh-Cre/tdTomato germ cell lineage tracing system;” see the “Sorting” section and Fig. 6 for more information. We genotype mice for Mvh-Cre and tdTomato by PCR, as described in “Materials and Methods.” Wild-type mice can also be used, but the purities obtained will be reduced (see Discussion for more details).
- WIN18,446, often sold under the name N,N′-Octamethylenebis(2,2-dichloroacetamide) or Fertilysin™: We source our WIN18,446 from MP Biomedicals (catalog number 02158050) or from Santa Cruz Biotechnology (catalog number sc-295819).
- Retinoic acid: It is most convenient to purchase this in small vials. We use Sigma Aldrich R2625-50mg.
- Carriers for WIN18,446 and RA suspensions: DMSO, PBS, and water. These should be sterile and of pharmaceutical grade.
- For injections: 1 ml syringes; 26-gauge needles, 3/8 inch or less in length. It is important to use a short needle of adequate diameter to reduce the risk that the WIN18,446 or RA suspension will clog the needle.

*Procedure:*

1) Begin with a single dam and her litter of pups, at p2. Before beginning dosing, we recommend removing any co-housed adult mice; this improves pup health and survival. If the litter is especially large (>6 pups for C57BL/6), we recommend euthanizing some of the female pups to reduce litter size. Also check that the dam is nesting well and that pups are healthy and of normal weight (for C57BL/6 mice, >1.5 g at p2). Developmental timing after synchronization is less consistent in low-weight pups.
2) Prepare WIN18,446 suspension fresh each day, directly before injection. For 1 ml, dissolve 5 mg of WIN18,446 in 25 µl of DMSO. Add 475 µl of sterile PBS and immediately shake to mix, generating a cloudy suspension. Always add PBS to WIN18,446/DMSO, rather than the reverse, to avoid precipitation of the WIN18,446.
3) Inject 10 µl per g body weight subcutaneously (over the shoulders) into male mice, once per day for 7 d. To reduce the risk of clogging, shake the WIN18,446 suspension well before filling the syringe, and load only ∼300 µl at a time. The first injection should be performed around noon; subsequent injections can be at any time of day. If schedule requires, a single day’s injection can be skipped, as long as injections are no more than 48 hours (h) apart. Note that pups sometimes develop a hairless patch on their neck from repeated injections; this is not harmful.
4) At p9 (day 8 of the protocol), check animal weight. If the animal is of healthy weight (>5 g in C57BL/6), it is ready for RA injection or tissue collection, as described below. In these normal-weight animals, germ cell development is expected to proceed with typical timing (Fig. 4A). If an animal is somewhat smaller (4-5 g), an extra 1-2 d of WIN18,446 can be administered to allow further weight gain before RA injection. These animals should be histologically staged with care. If animals have severely stunted growth (<4 g with an unhealthy appearance), it is best to euthanize them. If used, they should be carefully monitored, and germ cell development may be significantly slower than usual (Fig. 4B).
5) If the desired germ cell population consists of undifferentiated spermatogonia, euthanize the animal, dissect the testes and proceed to sorting and staging. Otherwise, prepare RA suspension for injection: first, make 100 mM RA stock (e.g. 50 mg RA + 1.664 ml DMSO). Then, mix 1.917 ml of water with 83 µl of RA stock. The result will be a yellow, opaque suspension. RA/DMSO stock can be stored for up to 7 d at −80°C. A number of cautions should be observed when preparing RA suspension. First, RA adheres to plastic, so use glass vials and pipettes whenever possible. Second, RA is oxygen-sensitive, so use the entire vial of RA powder at once when preparing the RA/DMSO stock. Finally, RA is light-sensitive, so store RA/DMSO stock in amber or foil-wrapped vials. Inject 10 µl of RA suspension per g of body weight, subcutaneously. Shake the RA suspension well before injection. Record the exact injection time, and animal weight at the time of injection. Be careful not to inject more than 10 µl per g of body weight, as too much RA can be toxic to pups and may stunt growth.
6) (Optional, but recommended) After RA injection, remove or euthanize any co-housed un-injected pups. RA-injected pups recover better if they do not need to compete with un-injected animals.
7) Monitor pups after injection, and choose the collection time-point based on the desired germ cell population. Developmental timing will have to be determined anew for each strain, and may vary somewhat from investigator to investigator even when working with the same strain. As a starting reference, our observed timings in normal-weight C57BL/6 animals are shown in Fig. 3 and Fig. 4A. If a pup appears unhealthy, or is not gaining weight appropriately, do not expect normal germ cell developmental timing (Fig. 4B). It may be best to euthanize the animal. At the time of collection, euthanize the animal, dissect out the testes, and proceed to sorting and staging.

Fig. 2 shows typical results of synchronization. In this experiment, development has been allowed to proceed for 9 d after the RA injection, in order to collect late leptotene spermatocytes. Sections from an unsynchronized control and a synchronized testis are stained for two meiotic markers: γH2AX, which marks double-strand breaks (Rogakou et al., 1998), and SYCP3, a component of the synaptonemal complex (Yuan et al., 2000). In the unsynchronized control, the tubules display heterogeneous staining patterns: a few tubules contain late leptotene spermatocytes, but most contain only germ cells at earlier steps of mitotic and meiotic development. In contrast, in the synchronized testis, each tubule displays the same staining pattern, indicating that late leptotene spermatocytes are the most developmentally advanced (and most prevalent) germ cells in >95% of tubules.

Figs. 3 and 4 show the enrichment of different germ cell populations at various time-points after RA restoration in C57BL/6 mice. Immediately prior to RA restoration (Fig. 3A), the only germ cells in the testis are undifferentiated spermatogonia (Fig. 1A). At 5.75 d after RA restoration, the most advanced germ cells in each tubule are type B spermatogonia; the testis also contains some undifferentiated spermatogonia (Fig. 3B). At 7 d after RA restoration, the most advanced germ cells are preleptotene spermatocytes that are positive for the meiotic initiation marker STRA8 (Baltus et al., 2006; Mark et al., 2015) (Fig. 3C). At 8, 10, and 12 d after RA restoration, the most advanced germ cells are progressively further into meiotic prophase (Fig. 3 D-F). We note that this timetable of germ cell development is more rapid than is usually observed in the adult, consistent with previous observations that pubertal spermatogenesis proceeds more rapidly than adult spermatogenesis (de Rooij and Russell, 2000; Kluin et al., 1982). From 7 d onward, a new generation of germ cells begins differentiating synchronously behind the meiotic cohort (Fig. 3 C-F) (Russell et al., 1990), increasing the total number of spermatogonia in the testis. In normal-weight animals, the timetable of synchronized germ cell development is quite consistent from animal to animal, and synchronization does not deteriorate significantly during these 12 d of development (Fig. 4A). We calculate that, throughout this period, germ cell development is synchronized to within a 24 h developmental window. This is a significant improvement over the unsynchronized juvenile testis—which has frequently been used as a less cellularly complex alternative to the adult testis: in the unsynchronized juvenile testis, the most- and least-advanced members of the first cohort of differentiating germ cells are separated by more than 3 d of development (Nebel et al., 1961; Russell et al., 1990).

### Step 2: Staging

To establish the timing of germ cell development after RA restoration, it is important to histologically examine (“stage”) a portion of one testis from each synchronized animal (Fig. 1B-2). By identifying when each step of germ cell development occurs, we can determine the typical timing of germ cell development in the mouse strain of interest. Even after the synchronization protocol is established in a given strain, it is still critical to stage every animal, for several reasons. First, developmental timing is only predictable in healthy, normal-weight animals; staging allows underweight or marginal animals to be used. Second, even in normal-weight animals, developmental timing can vary slightly (Fig. 4A); staging is thus essential for capturing short-lived cell populations. Finally, staging ensures that rare errors in synchronization are detected so that samples may be excluded from analysis.

We use half of one testis for staging, and the remainder for sorting. We stage and sort in parallel. In particular, we sort and process germ cells while the tissue for staging is being fixed. We then store sorted cells (or extracted RNA, protein, etc.) until staging is complete, whereupon we pool sorted cells from multiple animals and proceed with downstream studies.

The technique we use for preparing tissues is:

1) Cut one testis in half, using small dissection scissors to split the tunica albuginea. Fix the half testis and associated piece of tunica for 4 h in Bouin’s solution.
2) Embed tissue in paraffin, section (5 µm width), and stain with hematoxylin.

Then, for each tubule cross-section, determine its stage: that is, the most advanced (and most prevalent) step of germ cell development. Staging requires a light microscope with an oil objective and 100x magnification. Staging takes some effort to learn; two excellent references are Ahmed and de Rooij (2009) and Russell *et al.* (1990).

Some practical tips for learning to stage:

- By histology alone, leptotene is one of the easiest phases of germ cell development to identify; see the characteristic speckled hematoxylin staining in Fig. 3D.
- Preleptotene spermatocytes are also reasonably easy to identify by histology. For confirmation, sections can be stained with an antibody to the meiotic initiation marker STRA8, as described in “Materials and Methods” (Fig. 3C).
- Later phases of meiosis (zygotene, early/mid/late pachytene, diplotene) are more difficult to identify by histology, but their identity can be readily confirmed by meiotic spreads (de Boer et al., 2009; Peters et al., 1997).

Investigators can decide their own criteria for sample inclusion following staging. We include samples in which >90% of tubules contain germ cells at the desired step in their development. After synchronization and staging, testes are already substantially enriched for the germ cell population of interest, though of course somatic cells and spermatogonia are still present (Fig. 3). These synchronized and staged samples (“2S”) can in some cases be used directly for downstream experiments, but in most cases investigators will want to proceed to sorting (Fig. 1B-3).

### Step 3: Sorting

Sorting separates the desired germ cell population from contaminating cells, most of which are somatic, to produce high purities (Fig 1B-3). Because synchronization dramatically reduces the cellular complexity of the testis, and staging conclusively identifies those cells, relatively simple sorting strategies are sufficient to isolate the desired germ cells. We describe two sorting strategies (Fig. 5). The first, based simply on light scattering properties, gives moderately high purity, with no need for staining or genetic markers. The second, based on the Mvh-Cre/tdTomato germ cell lineage tracing system (Fig. 6), is more involved but yields ∼90% purity of multiple germ cell populations. Both strategies begin with a two-step digestion protocol (modified from Gaysinskaya et al., 2014) to reduce the testes to a single-cell suspension.

**Fig. 5.**
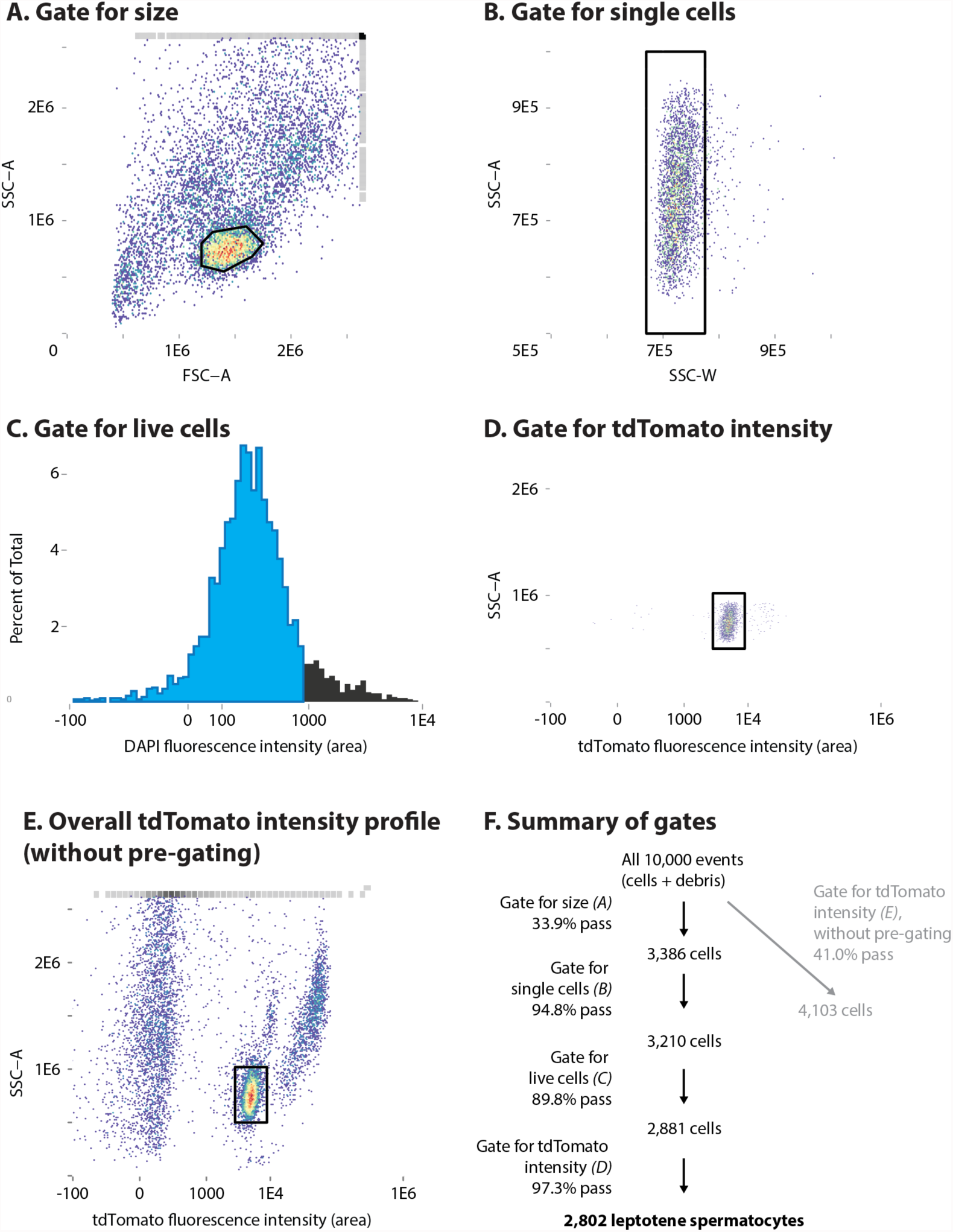
Gating for leptotene spermatocytes. (A) We first gate for the leptotene spermatocyte population by cell size, as assayed by forward and side light scatter (black outline). Red represents areas of highest density on the scatter plot. FSC-A: forward scatter-area; SSC-A: side scatter-area. (B) We next gate for single cells (black outline). Only cells that pass the size gate (A) are included in the scatter plot. SSC-W: side scatter-width. (C) We next gate for live (DAPI-negative) cells (blue). Only cells that pass the size and single-cell gates (A and B) are included in the histogram. (D) Finally, we optionally gate for tdTomato intensity (black outline). Only cells that pass the size, single-cell, and live-cell gates (A, B, and C) are included in the scatter plot. (E) Overall tdTomato fluorescence intensity profile. The scatter plot shows all data (without pre-gating for size, single cells, or live cells). The gate, for mid-range tdTomato fluorescence intensity (black outline), is the same as in (D); this population is primarily leptotene spermatocytes. The tdTomato-low population is primarily somatic cells; the tdTomato-high population is primarily spermatogonia (Fig. 6B). (F) Summary of the gates, and the number of cells that pass each gate. We present gating results on a subset of our data, encompassing 10,000 “events.” (Each event is a particle detected by the instrument, either a cell or a piece of non-cellular debris). Black part of flow chart shows the gates in (A, B, C, and D); gray part shows the gate in (E).

**Fig. 6.**
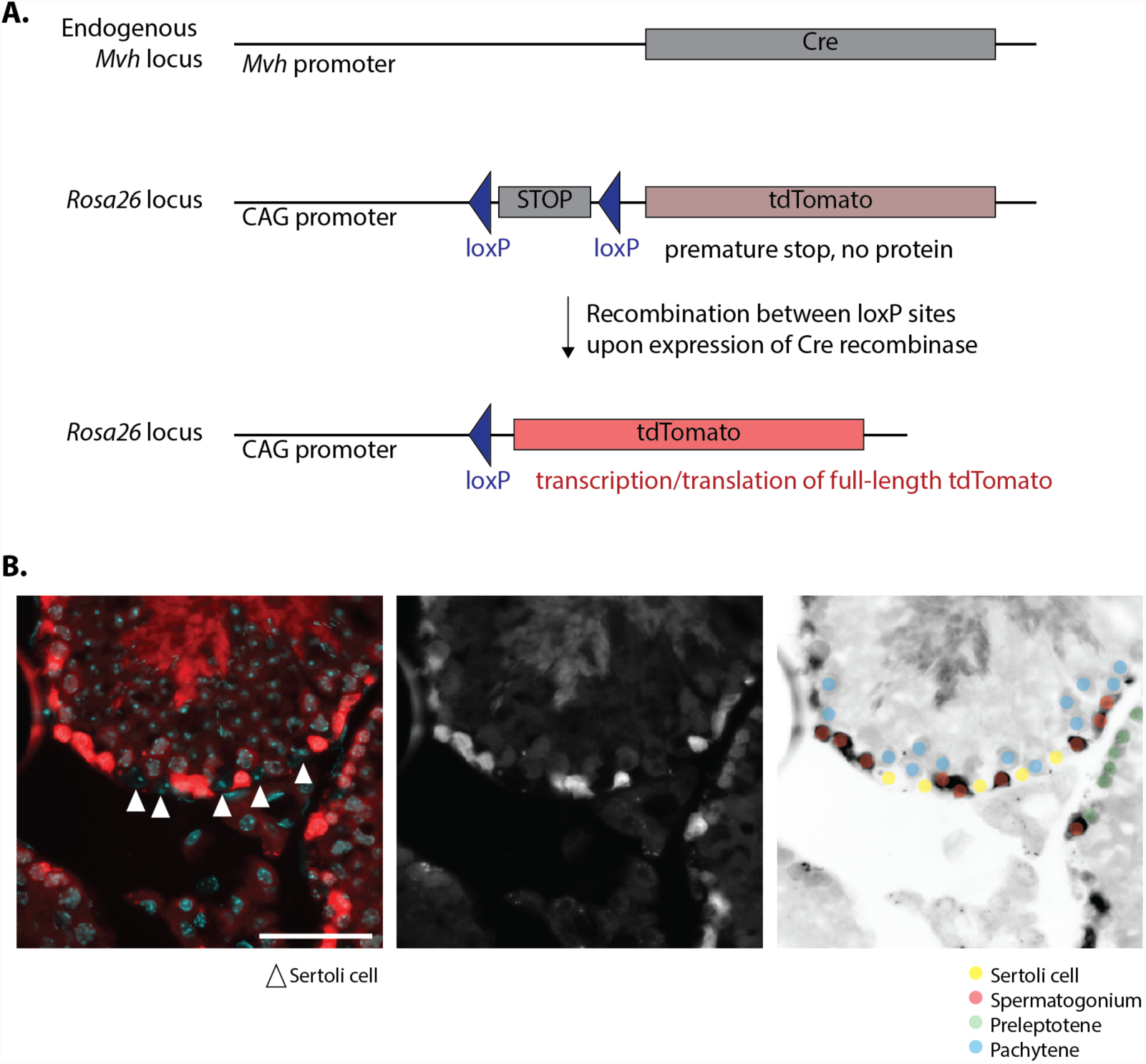
The Mvh-Cre/tdTomato germ cell lineage tracing system. (A) The Mvh-Cre/tdTomato lineage tracing system consists of two components. First, our mice are heterozygous for Mvh-Cre (Hu et al., 2013): Cre recombinase driven by the *Mvh* promoter, in the endogenous *Mvh* locus. (The other *Mvh* allele is wild-type.) Mvh-Cre has a similar expression pattern to wild-type *Mvh*; thus, all germ cells express Mvh-Cre at some point during their embryonic development. Mvh-Cre is not expressed outside the germline. Second, our mice are heterozygous for a loxP-STOP-loxP-tdTomato allele (Madisen et al., 2010), driven by the CAG promoter in the *Rosa26* locus. (The other *Rosa26* locus is unmodified.) In the absence of Cre, the STOP codon prevents translation of tdTomato. Upon exposure to Cre, the STOP codon is excised and tdTomato translation begins. These two alleles together function as a germ cell lineage tracing system; tdTomato protein is present in germ cell lineages that have activated the Mvh-Cre. (B) Pattern of tdTomato expression in the adult (unsynchronized) testis. Left panel shows a merge of tdTomato (red) and DAPI (cyan), with Sertoli cells indicated by white arrows; middle panel shows tdTomato alone (white). Right panel is an inverted version of the middle image, with different cell types marked with translucent dots: Sertoli cells (yellow), spermatogonia (red), preleptotene spermatocytes (green), and pachytene spermatocytes (blue). Scale bar, 50 µm.

*Materials and equipment:*

- Collagenase type I (we use Worthington, Cat # LS004196)
- 2.5% trypsin stock solution
- DNAse I, 6820 U/ml
- Hank’s balanced salt solution (HBSS)
- Serum, for quenching digestion (we use Hyclone cosmic calf serum)
- 40 µm filter
- DAPI (4′,6-diamidino-2-phenylindole) stock solution, 0.5 mg/ml in water
- FACS instrument. We use a FACSAria II instrument (BD Biosciences)

*Procedure*

1) Make fresh a collagenase solution (HBSS + 1 mg/ml collagenase + 1:1000 DNAse), and a collagenase/trypsin solution (HBSS + 1 mg/ml collagenase + 0.05% trypsin + 1:1000 DNAse). Warm both solutions to 35°C.
2) Remove the tunica albuginea from the collected testes and place the testes in 1 ml of collagenase solution in a 1.7 ml microtube.
3) Shake the tube vigorously by hand until the tubules begin to separate from each other. Then, slowly, gently shake in a horizontal position for 7 min at 35°C. (Shaking may be done manually or mechanically, and may be performed at room temperature rather than at 35°C if necessary.) Halfway through this shaking period, gently pipette the tubules up and down with a 1 ml pipette tip to assist in the tubule dispersion. By the end of the 7 min period, most tubules should appear thin and dispersed, though a few clumps may remain.
4a) (Standard protocol.) Place the tube in a vertical position and leave undisturbed at room temperature for 2 min, to allow tubules to settle to the bottom. Remove and discard the supernatant (which contains unwanted somatic cells from the testicular interstitium), leaving enough liquid to cover the tubules generously. Add 1 ml of collagenase/trypsin solution.
4b) (Alternate protocol to minimize tissue loss, for small quantities of tissue such as testes collected at 0 d after RA injection.) Do not remove supernatant. Instead, after tubule dispersion, add trypsin stock solution directly to the tubule/collagenase mixture, to achieve a final concentration of 0.05% trypsin.
5) Gently shake in a horizontal position for 20 min at 35°C. Every 5 min, gently pipette the tubules up and down to assist in digestion. After 10 min, add an additional 8 µl of 2.5% trypsin; if digestion solution is overly viscous, also add an additional 1 µl of DNAse I stock. At the end of this digestion period, the testis tubules should be reduced to a single-cell suspension.
6) Quench digestion with 400 µl of serum, mixing thoroughly by pipetting.
7) Wash, filter, and stain the testicular cells in preparation for sorting: spin down the cell suspension at room temperature for 5 min at 2000g, remove the supernatant, and re-suspend in 500 µl of ice-cold HBSS, making sure to thoroughly break up the cell pellet by repeated pipetting. Keep the cells on ice until ready to sort.
8) Immediately before sorting, pass the cell suspension through a 40 µm filter, and wash through with an equal volume of ice-cold HBSS. Add 2 µl of DAPI stock solution, to exclude dead cells during sorting.
9) Isolate germ cells by FACS. We use an 85 µm nozzle, and the gating strategy described below.

*Gating for germ cell subpopulations:* Fig. 5 shows our standard gating strategy, which gives high purity for meiotic and late mitotic cell populations, from type B spermatogonia through early pachytene spermatocytes. Specifically, Fig. 5 shows the gating results from a sort designed to enrich for leptotene spermatocytes. The input to this sort was a pair of synchronized testes, collected 8 d after RA restoration, and verified by staging to contain leptotene spermatocytes.

We first gate based on light scatter (forward and side scatter area), which largely assays cell size (Shapiro, 2003) (Fig. 5A). The majority of cells in this sample are leptotene spermatocytes (Fig. 3D), which are homogeneous in light scatter and can be easily gated. Next, we gate for single cells, using light scatter (Shapiro, 2003), and we gate for live cells, using DAPI fluorescence (Fig. 5B and Fig. 5C). For DAPI detection, we use a 375 nm UV laser and a 450 nm/520 nm bandpass filter. After these three gating steps, we are ready to sort a highly purified population of leptotene spermatocytes (Fig. 5F). We note that, because the target cell population is well separated from contaminating cells (Fig. 5A), we can use a fairly fast sorting rate without compromising sort efficiency: on the FACSAria II instrument, we use a flow rate of ∼60 µl/min.

To achieve greater purity, we use transgenic mice that carry a fluorescent marker of germ cells (Fig. 6). With this Mvh-Cre/tdTomato germ cell lineage tracing system, all germ cells in the postnatal testis express some level of the fluorescent protein tdTomato under the control of the CAG promoter (Fig. 6A). In somatic cells, tdTomato protein is absent, due to the presence of a STOP codon that is excised in germ cells. While the CAG promoter is generally thought to confer ubiquitous expression (Madisen et al., 2010), we observed that tdTomato fluorescence intensity in fact decreases gradually over the course of germ cell development, being brightest in early spermatogonia and only slightly above background in mid/late pachytene spermatocytes (Fig. 6B). For detection of tdTomato fluorescence, we use a 488 nm blue laser and a 585 nm/642 nm bandpass filter. Fig. 5E shows three distinct levels of tdTomato fluorescence, reflecting the three major cell populations in the synchronized testis: somatic cells (negative), leptotene spermatocytes (moderate), and spermatogonia (high). Combining tdTomato fluorescence with our previous gating strategy (Fig. 5D), we obtain a more precisely defined population of leptotene spermatocytes (Fig. 5F and Table 1). We find that a nearly identical gating strategy can be used for other late mitotic and early meiotic cell populations: STRA8-positive preleptotene spermatocytes (Fig. 7A-C), and zygotene spermatocytes (Fig. 7D-F). The Mvh-Cre/tdTomato lineage tracing system is helpful but not necessary for sorting these populations. In contrast, our experience indicates that the Mvh-Cre/tdTomato lineage tracing system is necessary for the sorting of undifferentiated spermatogonia (Fig. 8A-D) and is helpful for sorting early differentiating spermatogonia, because spermatogonia are somewhat heterogeneous in size (Fig. 8E-F) and because, at such early time-points, somatic cells outnumber germ cells (Fig. 2A).

**Table 1.**
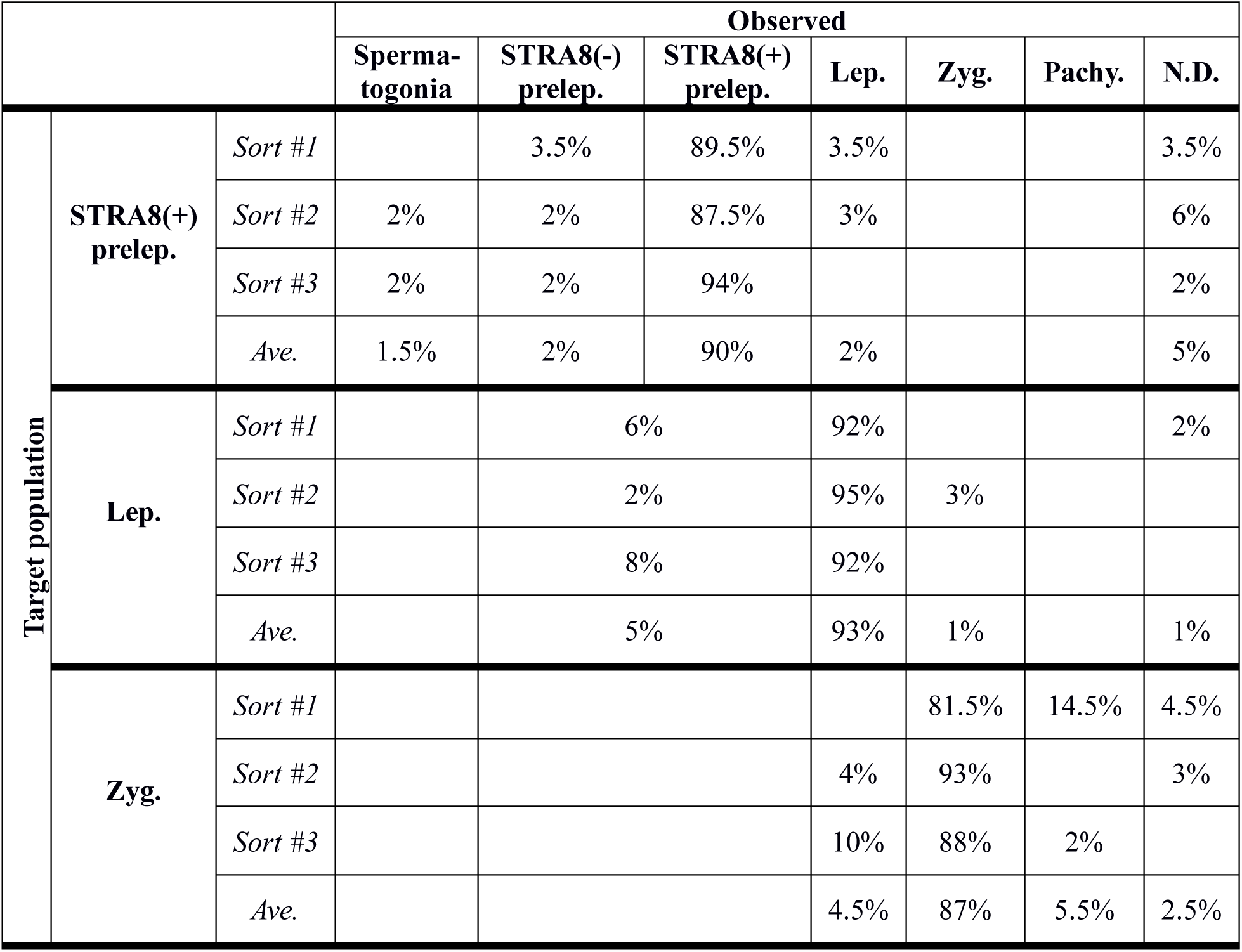
Purities of sorted cell populations. For each target population (STRA8-positive preleptotene, leptotene, or zygotene spermatocytes), three sorts were performed, each from a single synchronized and staged animal. At least 50 randomly selected cells were assessed for each animal. Purities were assessed by immunostaining (Fig. 9). For the leptotene and zygotene sorts, STRA8-positive and STRA8-negative preleptotene spermatocytes were not distinguished. Purities are given rounded to the nearest 0.5%. Prelep., Lep., Zyg., Pachy.: preleptotene, leptotene, zygotene, and pachytene spermatocytes. N.D.: not determined.

**Fig. 7.**
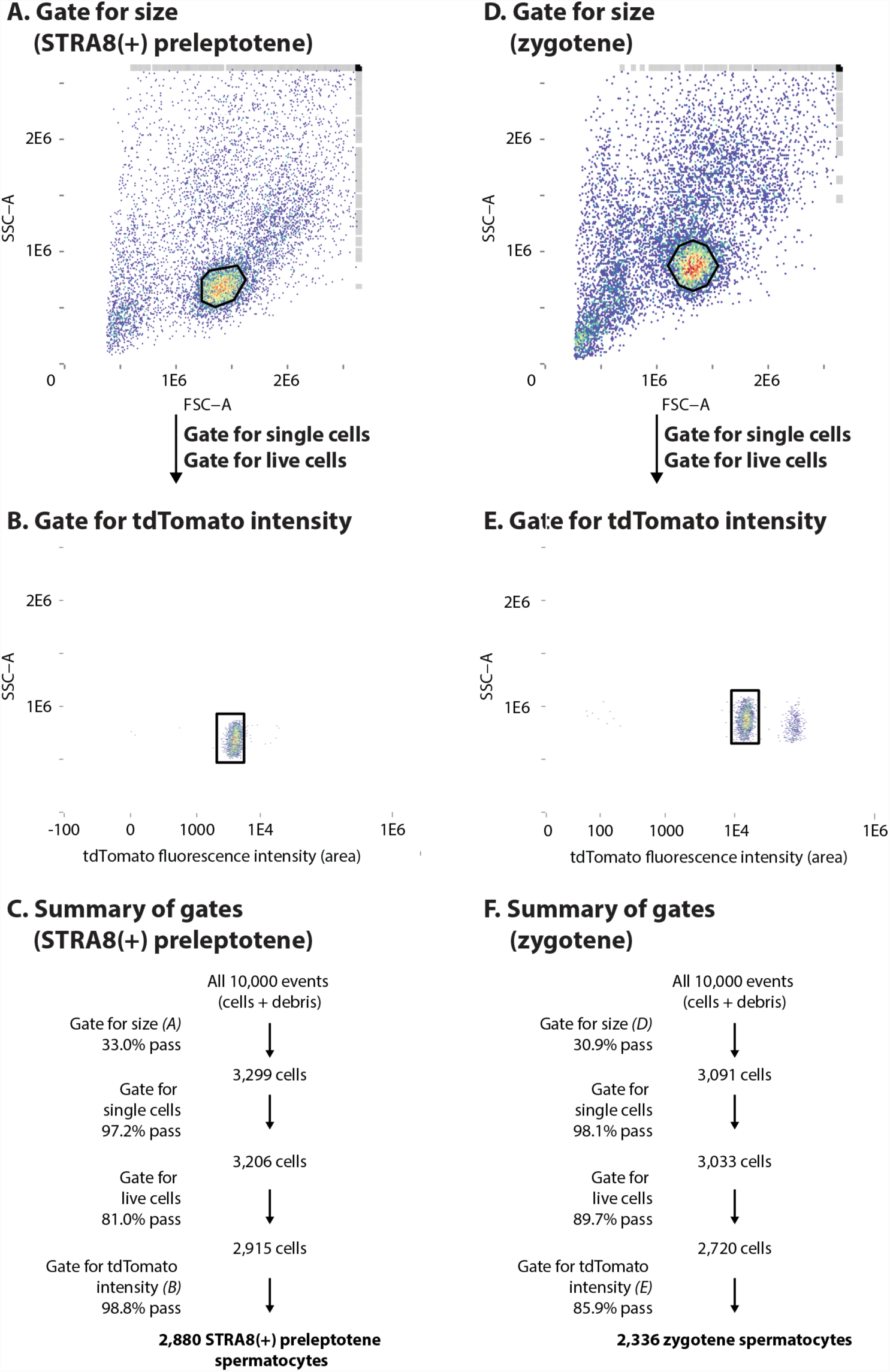
Gating for STRA8-positive preleptotene spermatocytes and for zygotene spermatocytes. (A and D) We first gate for the STRA8-postive preleptotene (A) and zygotene (D) spermatocyte populations by cell size (black outlines). Red represents areas of highest density on the scatter plot. FSCA: forward scatter-area; SSC-A: side scatter-area. Following the size gate, we gate for single cells and for live cells as in Fig. 5 B and C. (B and E) Next, we optionally gate for tdTomato intensity (black outlines) to isolate STRA8-positive preleptotene (B) and zygotene (E) spermatocytes. Only cells that pass the size, single-cell, and live-cell gates are included in the scatter plot. (C and F) Summary of the gates, and the percentage of cells that pass each gate.

**Fig. 8.**
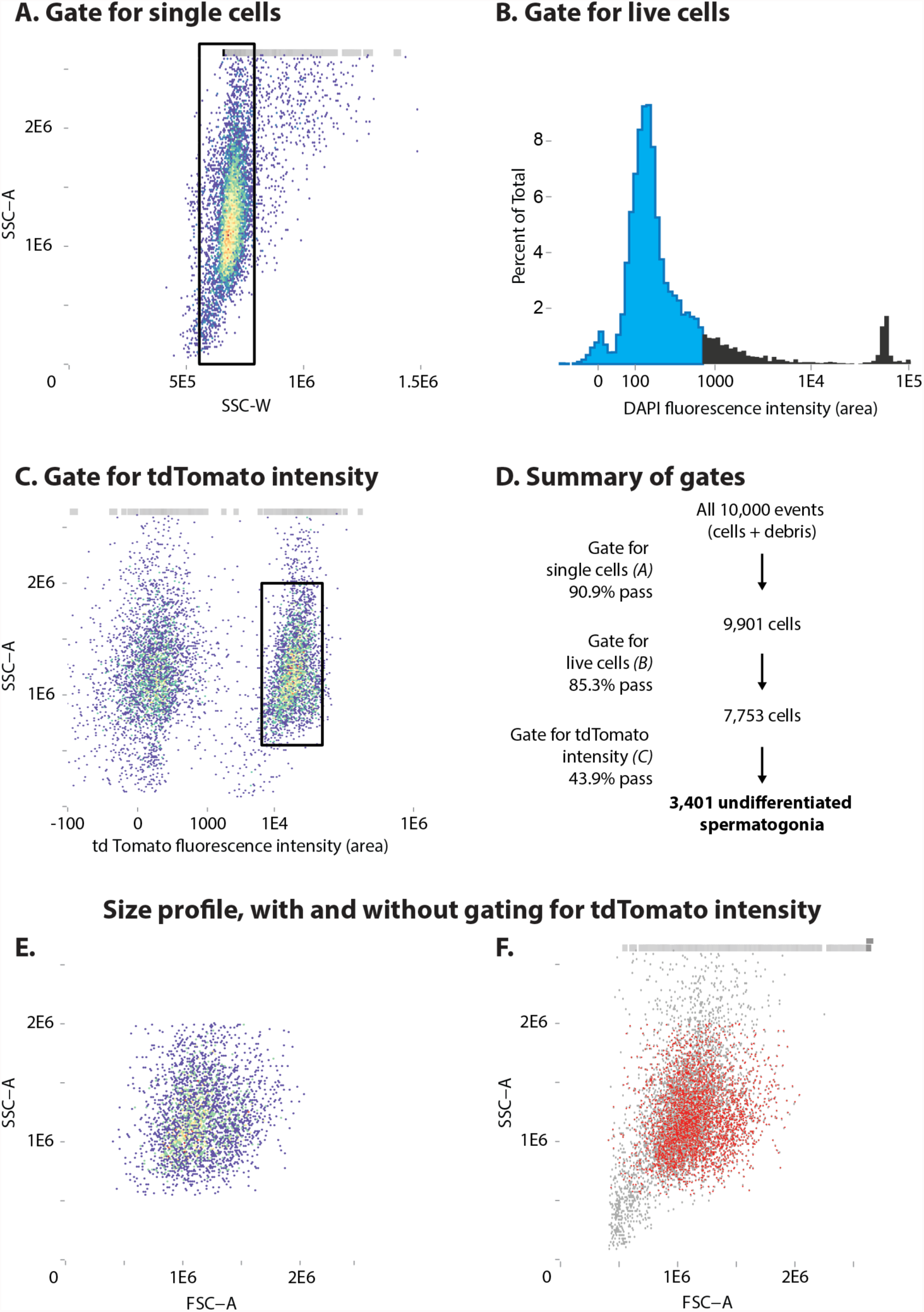
Gating for undifferentiated spermatogonia. (A) We first gate for single cells (black outline). Red represents areas of highest density on the scatter plot. SSC-A: side scatter-area; SSC-W: side scatter-width. (B) We next gate for live (DAPI-negative) cells (blue). Only cells that pass the single-cell gate (A) are included in the histogram. (C) Finally, we gate for tdTomato intensity (black outline). Only cells that pass the single-cell and live-cell gates (A and B) are included in the scatter plot. (D) Summary of the gates, and the percentage of cells that pass each gate. (E) Light scatter (size) profile of putative undifferentiated spermatogonia. Only cells that pass the single-cell, live-cell, and tdTomato-intensity gates (A*-*C) are included in the scatter plot. Putative undifferentiated spermatogonia do not cluster tightly on the scatter plot, indicating a fairly wide range of sizes. We believe that this wide range of sizes is observed because the cell cycle of undifferentiated spermatogonia is not subject to synchronization (van Pelt et al., 1995). FSC-A: forward scatter-area; SSC-A: side scatter-area. (F) Light scatter profile of putative undifferentiated spermatogonia (gated for single cells, live cells, and tdTomato intensity, from (E), in red), overlaid on overall light scatter profile (all data, ungated, in gray). This shows that the size range of putative undifferentiated spermatogonia overlaps with that of somatic cells of the testis.

### High purities obtained by synchronizing, staging, and sorting

We have used the 3S approach to isolate, at high purities, germ cell subpopulations ranging from undifferentiated spermatogonia to early pachytene spermatocytes. As a representative test of the method, we have isolated and performed detailed purity assessments for three types of spermatocytes that had previously proved difficult to separate: late (STRA8-positive) preleptotene, leptotene, and zygotene. These encompass meiotic initiation in preleptotene, followed by key events of meiotic prophase: DNA double-strand break formation in leptotene, and break repair and homologous chromosome synapsis beginning in zygotene.

To enrich for these three phases of meiosis, we synchronize germ cell development and collect testes at 7, 8, and 10 d after RA restoration (Fig. 3 C, D, and E). We stage each sample, requiring that at least 90% of tubules contain the target cell population; we discard any samples that did not meet this criterion. We then sort and perform purity assessments on the sorted cells (Table 1). To assess purity, we immunostain fixed cells (for preleptotene samples) or meiotic chromosome spreads (for leptotene and zygotene samples). We distinguish preleptotene, leptotene, and zygotene spermatocytes by staining for the meiotic initiation marker STRA8 and for the synaptonemal complex proteins SYCP3 (part of the lateral element) and SYCP1 (part of the axial element), using the criteria of Morelli *et al.* (2008) and Gaysinskaya *et al.* (2014). Representative immunostained cells and spreads are shown in Fig. 9.

**Fig. 9.**
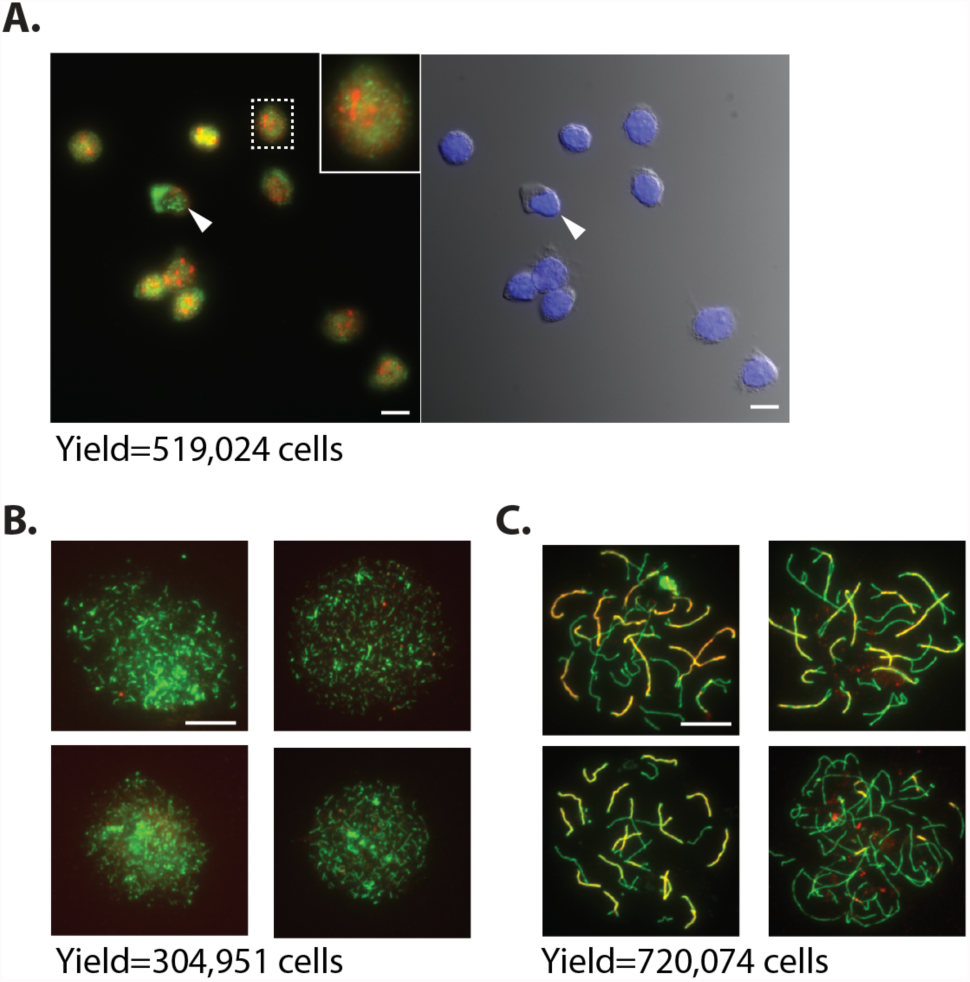
Immunostaining to determine purity of sorted spermatocyte populations. (A) Spermatocytes, fixed and immunostained, from a representative sort designed to enrich for STRA8-positive preleptotene spermatocytes. (Left and right images show the same field of cells). All but one of the cells can be clearly identified as STRA8-positive preleptotene spermatocytes, with the diffuse and punctate SYCP3 staining patterns characteristic of preleptotene (Gaysinskaya et al., 2014). One cell (white arrow) is STRA8-positive but SYCP3-negative with unusual morphology. Scale bars, 5 µm. Left: Immunostaining for STRA8 (green) and SYCP3 (red). Inset enlarges the boxed region. Right: Counterstaining for DAPI (blue), with differential interference-contrast shown in grayscale. (B and C) Meiotic spreads, from representative sorts designed to enrich for leptotene (B) and zygotene (C) spermatocytes. Immunostaining for SYCP3 (green) and SYCP1 (red). All cells in (B) can be clearly identified as leptotene spermatocytes, with stretches of SYCP3 along parts of the chromosome axes and absent or diffuse SYCP1 staining. All cells in (C) can be clearly identified as zygotene spermatocytes, with SYCP3 along the entire chromosome lengths and stretches of SYCP1 (Gaysinskaya et al., 2014; Morelli et al., 2008). Scale bars, 15 µm. In (A, B, and C), below the image, we indicate the number of cells yielded by the representative sort.

We obtain high purity (∼90%) for preleptotene, leptotene, and zygotene cells (Table 1), demonstrating that we can consistently separate these spermatocyte populations. When we compare these results to published data from unit gravity sedimentation and from DNA staining/FACS (Table 2) (Bellvé, 1993; Gaysinskaya et al., 2014), we find that the purities obtained with 3S represent a substantial improvement over previous methods. In particular, 3S is able to separate leptotene and zygotene spermatocytes with higher purity than was previously possible. Moreover, we are able to specifically isolate the subset of preleptotene spermatocytes that had initiated meiosis, as assessed by staining for the marker STRA8 (Fig. 9A). To our knowledge, this is the first time that STRA8-positive spermatocytes have been specifically isolated at high purity: the unperturbed testis contains a mixture of STRA8-positive and -negative preleptotene spermatocytes, which cannot be separated by unit gravity sedimentation or DNA staining/FACS. Separating STRA8-positive from STRA8-negative preleptotene spermatocytes should enable a better understanding of meiotic initiation and the earliest events of meiotic prophase.

**Table 2.**
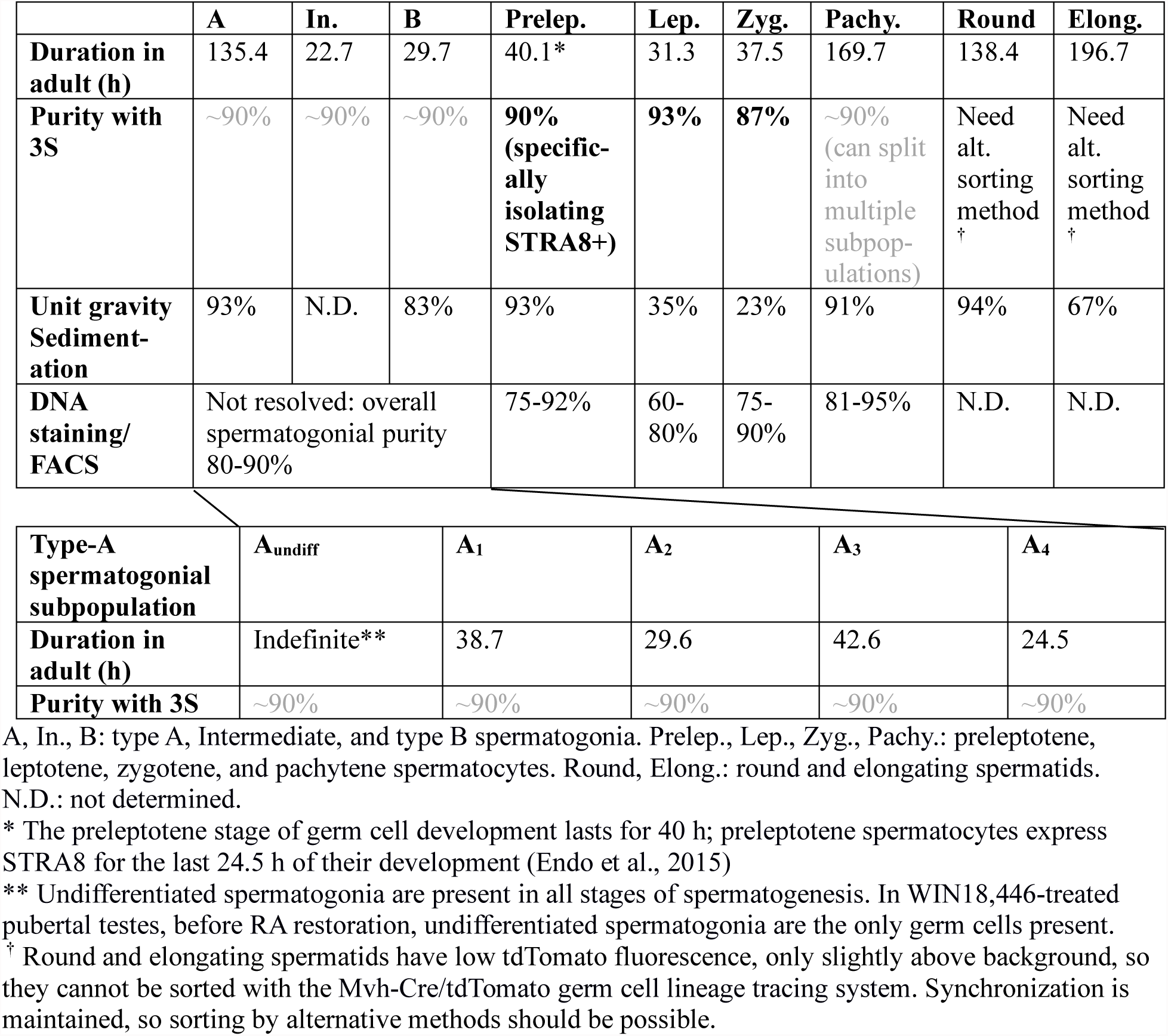
Comparison of purities obtainable with 3S, unit gravity sedimentation, and DNA staining/FACS. Purities obtainable with unit gravity sedimentation and DNA staining/FACS are from (Bellvé, 1993) and (Gaysinskaya et al., 2014) respectively. For DNA staining/FACS, all cell populations were sorted from adult testes. For unit gravity sedimentation, some populations were sorted from juvenile testis. Bottom table indicates that 3S can split Type-A spermatogonia into multiple subpopulations, which is not possible with the other two methods. For 3S, percentages in bold represent measured purities (Table 1); percentages in gray represent estimated purities for populations that we have not yet tested. Durations of each step of development are calculated based on (Ahmed and de Rooij, 2009; Oakberg, 1956).

Based on these findings, we believe that 3S can be used to isolate a wide range of mitotic and meiotic germ cell populations, at high purity. We have demonstrated that synchronization is maintained throughout both mitotic and meiotic germ cell development; throughout these processes, germ cell development is synchronized within a 24 h window. We have also found that 3S produces consistent purities of ∼90% for the three tested steps of germ cell development. Thus, we expect that 3S can be used to isolate, at approximately 90% purity, any step of development that lasts for 24 h or more, from the undifferentiated spermatogonia until the loss of tdTomato fluorescence in early/mid pachytene (Table 2). Thus, 3S should be able to separate the different populations of spermatogonia––undifferentiated, A_1_, A_2_, A_3_, A_4_, Intermediate, and B––in addition to different populations of spermatocytes––preleptotene, leptotene, zygotene, and early and mid pachytene.

## DISCUSSION

Our 3S approach yields higher purity and finer-grained separation of germ cell subpopulations than was achievable with the previous state-of-the-art methods: unit gravity sedimentation (Bellvé, 1993) and DNA staining/FACS (Gaysinskaya et al., 2014) (Table 2). In the mitotic phase, 3S enables, for the first time, systematic separation of the steps of spermatogonial development. We anticipate that this will lead to better understanding of spermatogonial self-renewal, proliferation, and preparation for meiosis. In the meiotic phase, 3S yields improved purity of preleptotene, leptotene, and zygotene spermatocytes; enables specific isolation of preleptotene spermatocytes that are positive for the meiotic initiation marker STRA8; and allows separation of multiple subpopulations of pachytene spermatocytes. This fine-grained separation of meiotic cell populations should improve understanding of the molecular events of meiotic initiation and early prophase—fundamental processes that occur in organisms from yeast to man.

In addition to providing precise germ cell populations at higher purity, the 3S protocol has several practical advantages over other methods. First, one uses the same basic protocol to sort many different cell populations, from undifferentiated spermatogonia to mid-meiotic prophase. To obtain different cell populations, one simply changes the amount of time that germ cells are allowed to develop after RA restoration. Reflecting this consistency, the purities obtained for preleptotene, leptotene, and zygotene spermatocytes are quite similar. This is in contrast to other sorting methods, where the purity depends on the cell populations being sorted, and where investigators must optimize the protocol anew for each population.

Second, cells are treated gently in 3S. Because synchronization is achieved *in vivo*, the testis is subjected to minimal handling after dissection. It does not need to be stained with cytotoxic dyes (Durand and Olive, 1982), and sorting can be conducted quickly. Because most cells are alive at the time of sorting (Fig. 5C and Fig. 8B), the transcriptional and epigenetic state of the cells is likely to be relatively unperturbed after sorting.

Third, 3S gives high yields. For the isolation of spermatocytes, we obtained ∼500,000 cells from each pair of testes, in 15-20 min of sorting time (Fig. 9). This high yield is largely due to synchronization. In the unperturbed adult testis, the percentage of germ cells in any given phase of meiosis is low (Gaysinskaya et al., 2014); synchronization increases this percentage by a factor of 10 or more (Russell et al., 1990), allowing rapid sorting. The yield from a single animal, or in some cases several pooled animals, is sufficient for many genome-wide assays, including deep transcriptional profiling and biochemical assays, including ChIP-seq for epigenetic marks and transcription factor binding, which are not suitable for single-cell approaches.

Finally, 3S gives excellent results without specialized equipment and supplies. The Mvh-Cre/tdTomato germ cell lineage tracing system gives the best purities in our hands. However, this transgenic system is not required: when sorting spermatocyte populations, simply selecting for light scatter properties can provide purities of >75% (Fig. 5 and Fig. 7). If a FACS instrument is not available, synchronization and staging without sorting (“2S”) can be used (Figs. 3 and 4). We note that 2S represents a dramatic improvement over the commonly used “first wave of spermatogenesis,” which attempts to enrich for particular cell populations by timed collection of unsynchronized pubertal testes (Nebel et al., 1961).

In the future, we anticipate that 3S can be extended to allow isolation of late meiotic and post-meiotic populations (round and elongating spermatids). Synchronization is maintained throughout post-meiotic spermatid development (Hogarth et al., 2013), so the main obstacle is identifying sorting methods suitable to these populations. Since tdTomato fluorescence intensity diminishes over the course of germ cell development, our Mvh-Cre/tdTomato lineage tracing system cannot be used to sort germ cells after mid-pachytene of meiosis, and by mid-pachytene the cell composition of the testis has become too complex to sort by light scattering properties alone (Fig. 3F). However, a variety of alternative sorting methods are available to isolate late meiotic and post-meiotic cell populations. The simplest is combining synchronization and staging with a DNA staining/FACS approach. DNA staining excels at separating germ cells by ploidy, so it is a natural approach for isolating haploid spermatids, and can be used to isolate 4N spermatocytes as well (Gaysinskaya et al., 2014). This extension of 3S would enable greater understanding of the meiotic divisions, and of the cellular remodeling that occurs during haploid differentiation.

We believe that 3S will enable investigators to pursue biological questions that were previously difficult or impossible to experimentally address. First, 3S could enable better understanding of the changes that occur during the long mitotic phase of spermatogenesis. Over the course of these transit-amplifying divisions, spermatogonia lose expression of pluripotency genes (Chakraborty et al., 2014) and gain competence for meiosis. By isolating different spermatogonial types (A_1_ through B) along this developmental progression, one could begin to understand these changes at a molecular and biochemical level. Second, by enabling separation of STRA8-negative and -positive preleptotene spermatocytes, 3S could reveal the gene expression changes that occur immediately before and after meiotic initiation, leading to better understanding of the regulatory network governing meiotic initiation. Third, by enabling isolation of large numbers of preleptotene, leptotene, and zygotene spermatocytes, 3S opens the door to more detailed genome-wide and biochemical characterizations of these phases of meiosis. For example, one could profile how meiotic factors involved in double-strand break formation and homologous recombination (PRDM9, SPO11, and DMC1) are bound to DNA in each of these phases. To date, the most detailed studies of progression through meiotic prophase have been conducted in yeast, in part due to the ease of synchronizing meiosis and sporulation; 3S could enable a clearer understanding of how meiotic prophase differs between mammals and lower eukaryotes (Chu et al., 1998; Hayase et al., 2004; Hunter, 2003; Panizza et al., 2011; van Werven and Amon, 2011). Finally, over both the mitotic and meiotic phases, 3S will aid in characterizing the epigenetic state of germ cells. DNA and histone methylation states are known to change during the course of spermatogenesis, but the precise dynamics and control of these changes are not understood. By using 3S to epigenetically profile precise steps of germ cell development, one could determine how imprinting and other epigenetic marks associated with normal embryonic development are established and maintained in germ cells, and how the epigenetic state of germ cells correlates, transgenerationally, with the epigenetic state and phenotype of offspring (Arnaud, 2010; Lesch et al., 2013; Sasaki and Matsui, 2008).

More broadly, the mammalian testis offers a potentially powerful model for studying a wide variety of biological processes, including cell proliferation and lineage commitment (Oatley and Brinster, 2008), the cell cycle, cellular remodeling (Russell et al., 1990), and alternative splicing and polyadenylation (Liu et al., 2007; Soumillon et al., 2013). However, testis biology can be daunting for the non-expert, due to the complex cell composition of the testis. 2S and 3S provide a straightforward means to reduce this complexity, and they do not require specialized equipment or large amounts of starting material. 2S and 3S could be applied by investigators across many subfields of modern biology and biochemistry; we hope that these techniques will encourage wider adoption of the testis as a model system.

## MATERIALS AND METHODS

### Mice

Three stains of mice were used: wild-type (BL/6NtacfBR); tdTomato (with a loxP-STOP-loxP-tdTomato construct in the *Rosa26* locus, strain B6.Cg-Gt(ROSA)26Sor^tm14(CAG-tdTomato)Hze^/J from The Jackson Laboratory) (Madisen et al., 2010); and Mvh-Cre (with a Cre-mOrange fusion protein, driven by the endogenous *Mvh* promoter) (Hu et al., 2013). To generate pups carrying the Mvh-Cre/tdTomato germ cell lineage tracing system, generally females homozygous for Mvh-Cre were crossed to males homozygous for tdTomato; however, in some cases animals heterozygous for Mvh-Cre or tdTomato were used for breeding. (Note that females homozygous for Mvh-Cre are fully fertile, while males homozygous for Mvh-Cre are sterile.) All animals were genotyped by PCR for Mvh-Cre and tdTomato before testis collection; genotyping for tdTomato was performed using the protocol provided by The Jackson Laboratory, and genotyping for Mvh-Cre was performed as previously described (Hu et al., 2013). All experiments involving mice were approved by the Committee on Animal Care at the Massachusetts Institute of Technology.

### Immunostaining of testis sections

Bouin’s-fixed, paraffin-embedded sections from synchronized testes were prepared as described in Results. Slides were dewaxed and rehydrated with a xylene and ethanol series. Antigen retrieval was performed by heating in a 10 mM sodium citrate buffer (pH 6.0). For STRA8 immunostaining, slides were blocked with 2.5% horse serum and 5% BSA for 90 min and incubated for 60 min at room temperature with primary antibody: 1:500 anti-STRA8 (Abcam ab49505, rabbit polyclonal antibody). Detection was colorimetric; slides were washed, incubated with ImmPRESS anti-rabbit IgG detection reagent (Vector Laboratories) for 30 min, and developed using a DAB substrate kit (Vector Laboratories). Slides were then counterstained with Mayer’s hematoxylin, dehydrated, and mounted with Permount (Fisher Scientific). For SYCP3 and γH2AX immunostaining, slides were blocked with 2.5% donkey serum for 30 min and incubated overnight at 4°C with primary antibodies: 1:250 anti-SYCP3 (Santa Cruz Biotechnology sc-74569, mouse monoclonal) and 1:150 anti-γH2AX (Pierce MA5-15130, rabbit monoclonal). Detection was fluorescent. Slides were washed and incubated for 60 min with 1:200 secondary antibodies (Jackson ImmunoResearch): Cy5-conjugated anti-rabbit IgG and rhodamine red X-conjugated anti-mouse IgG. Slides were mounted using Vectashield with DAPI (Vector Laboratories) and sealed with nail polish.

### Fixing and staining intact sorted cells

After sorting, cells were fixed for 15 min in 4% paraformaldehyde (PFA) and then washed twice in PBS. After fixing, cells were re-suspended in PBS and stored at 4°C. Staining was performed at room temperature. Cells were settled onto poly(L) lysine coated coverslips, permeabilized for 10 min by incubation in 0.25% Triton X-100, and blocked with 2.5% donkey serum and 1% BSA for 90 min. The coverslips were incubated for 60 min with primary antibodies: 1:500 anti-STRA8 and 1:100 anti-SYCP3 (see above). Coverslips were then washed with PBS and incubated for 60 min with 1:250 secondary antibodies (Jackson ImmunoResearch): Cy5-conjugated anti-rabbit IgG and FITC-conjugated anti-mouse IgG. Coverslips were mounted and sealed as above.

### Making and immunostaining meiotic chromosome spreads

Meiotic spreads were made following the protocol of (Peters et al., 1997). Briefly, after sorting, cells were re-suspended in hypotonic extraction buffer, and then in 100 mM sucrose. The cell/sucrose suspension was then pipetted onto glass slides, which had been coated in a thin layer of fixative (1% PFA and 0.15% Triton X-100). Slides were dried slowly in a humid chamber at room temperature. To stain, slides were washed with PBS and blocked with 3% BSA and 1% donkey serum for 30 min. Slides were then incubated with primary antibody overnight at 4°C: 1:250 anti-SYCP3 (see above) and 1:250 anti-SYCP1 (Abcam ab15090, rabbit polyclonal). Spreads were then washed with PBS and incubated for 60 min with 1:250 secondary antibodies (Jackson ImmunoResearch): Cy5 conjugated anti-rabbit IgG and rhodamine red X-conjugated anti-mouse IgG. Finally, spreads were washed with PBS, incubated with 0.02 µg/ml DAPI for 10 min, washed with water, mounted with Vectashield (Vector Laboratories), and sealed with nail polish.

### Data analysis

FACS data were analyzed with R, using the Bioconductor libraries flowCore and flowViz. Immunostaining images were processed with Fiji (Schindelin et al., 2012).

## ACKNOWLEDGEMENTS

We thank H. Christensen and M. Kojima for helpful discussions and assistance with synchronization; T. Endo for assistance with histological staging; the Whitehead Flow Cytometry Core (C. Johns, C. Zollo, and P. Wisniewski) for performing the sorting; J. Hughes for assistance with manuscript preparation and editing; and D.W. Bellott, H. Christensen, M. Kojima, and P. K. Nicholls for critical reading of the manuscript. Supported by the Howard Hughes Medical Institute and NIH Pre-Doctoral Training Grant T32GM007287.

